# Survey and Evaluation of Applied Containerization Practices in Bioinformatics

**DOI:** 10.64898/2026.07.17.739198

**Authors:** VP Nagraj, Stephen D Turner, Neal Magee

## Abstract

Containerization enables portable, reproducible, and scalable scientific computing. However, container development, documentation, and deployment practices can vary widely, even within a single domain. Understanding patterns of how containers are implemented in real-world settings can inform community guidelines, quality scoring rubrics, and systems to automate software review. Focusing on bioinformatics as an exemplar domain, we surveyed published software tools to find applied containerization examples. After identifying >250 tools to review, we annotated metadata regarding version control, asset provenance, and general container image health and tested the ability to build and pull the container images in a purpose-built evaluation platform (https://github.com/vpnagraj/socr8s). The majority of images tested did not build successfully in our environment. Of the tool characteristics we tracked, the strongest predictor of build success was version control activity. Tools with commits in the preceding two years were roughly twice as likely to build. Deeper assessment of failures identified a variety of issues, including missing assets and broken dependency chains. While many tools had published images accessible in open registries, 24% could neither be built nor pulled. Images tended to be large, with median size of 1.57GB for those that were able to be built. Where image specification files were available, we found that more advanced container orchestration and build techniques were uncommon. Our results highlight areas of improvement in how containerized tools are built and maintained in bioinformatics and scientific computing software in general.

## Background

Container technology enables portable, reproducible, and scalable computing practices. When tools and applications are packaged as a container image, the dependent software, configuration, and instructions for execution are defined in layers that are frozen at build time. Once instantiated, the container will execute as instructed on the host machine, using the specified dependencies, source code, and relevant built-in assets. Containerization confers particular benefits for scientific computing use-cases, which often require significant computational power and reliance on software that is openly available yet prone to breaking changes over time.

Given these advantages, containers have become critical to scientific computing workflows in numerous disciplines. Container technology has been adapted to meet research computing paradigms (e.g., shared high-performance computing clusters) via tools like Apptainer (formerly Singularity) (Kurtzer, Sochat, and Bauer 2017; Sochat, Prybol, and Kurtzer 2017). Researchers have proposed generalized guidelines for developing and implementing containerized tools (Moreau and Wiebels 2026; Nüst et al. 2020; Alser et al. 2024; Gruening et al. 2019). Some research teams have offered domain-specific perspectives that convey particular advantages of containerization practices. For instance, in quantitative psychology there are multiple publications that highlight the benefits of containers for reproducible science (Clyburne-Sherin, Fei, and Green 2019; Wiebels and Moreau 2021). Containerization has been presented as an efficient means to translate research computing tools into operational pipelines (Moreau, Wiebels, and Boettiger 2023). For example, one review describes applications of bioinformatics containers for genomic surveillance activities in public health laboratory settings (Florek et al. 2025). Despite the preponderance of published examples, tutorials, and informal guidelines for containerized development, studies of how container practices manifest in real-world scenarios have been limited. With the exception of one broad empirical study of the reproducibility of software using Docker (Malka, Zacchiroli, and Zimmermann 2026), there has been little evidence as to how, why, and to what effect containerization techniques are applied in computational science.

To better understand how containers are developed and deployed in scientific computing contexts, we set out to identify and review source code for a selection of recently published containerized bioinformatics tools. We chose to restrict our study to bioinformatics because containers are used for computational biology applications (Jackman et al. 2019; Kadri et al. 2022; Bai et al. 2021) and bioinformatics is a diverse discipline in terms of software tools and computational techniques used (Altschul et al. 2013; Mangul et al. 2019). The field also has a variety of publication outlets for “software papers” and therefore more opportunities to find published examples to review.

At the outset, we expected to find a wide range of approaches to building and deploying bioinformatics containers. Documenting and analyzing the techniques used in practice can help guide community standards, establish metrics for evaluating the rigor of containerized tools, and motivate software for auditing container quality. To that end, our study had three objectives:

1. Characterize how bioinformatics tools are containerized in practice
2. Identify the characteristics associated with whether a containerized tool can be successfully rebuilt or pulled
3. Create and release developer-facing tooling to support the study and improvement of containerized bioinformatics software

## Methods

### Literature search

To begin our study of published containerized tools, we first conducted a literature review. Given the bioinformatics focus, we considered PubMed as the repository of record for relevant publications. We developed an automated query of articles describing containerized bioinformatics software. Our query included terms to match bioinformatics use-cases using a range of possible synonyms with Medical Subject Headings (MeSH) in title, abstract, and body searches. In addition to the bioinformatics scope, we included a broad set of terms related to containerization. As a final step, to ensure that we captured the most current practices we set a date range for the published articles restricted from January 2021 to December 2025. We executed the query in late December 2025 using the NCBI Entrez Programming Utilities (E-utilities) API via the rentrez R package (Winter 2017) to search PubMed.

The E-Utilities API includes publication title, PubMed ID, abstract, authors, journal, and year published for all returned records. We used this information to manually audit the initial search results to confirm that the article described a containerized bioinformatics tool. We also assessed the scope of each article to determine if it was a software paper, experimental study, meta-analysis, review, protocol, or other type of manuscript. Only papers that describe novel implementations of containerized tools (either as a dedicated software paper or as a method in an experimental study) were retained for further review. Results from preprint servers (e.g., Biorxiv) were excluded. If the same tool was described in multiple papers, we retained only the most recent.

Using the screened results, we accessed the full text of each paper to retrieve links to source code and documentation for the containerized tool (e.g., GitHub repository) for additional screening. Tools that lacked links to source code or had no prebuilt images hosted on open repositories were excluded. Any tools that solely used off-the-shelf container images in pipelines, used containers only as a means to distribute data, or were provided as evidence of reproducibility for a specific analysis (i.e., bespoke images to run a pipeline on a specific dataset) were not included in further analysis. Tools explicitly marked as deprecated or abandoned were also excluded.

The initial PubMed query strategy and tool review protocol is illustrated in Figure 1.

**Figure 1:**
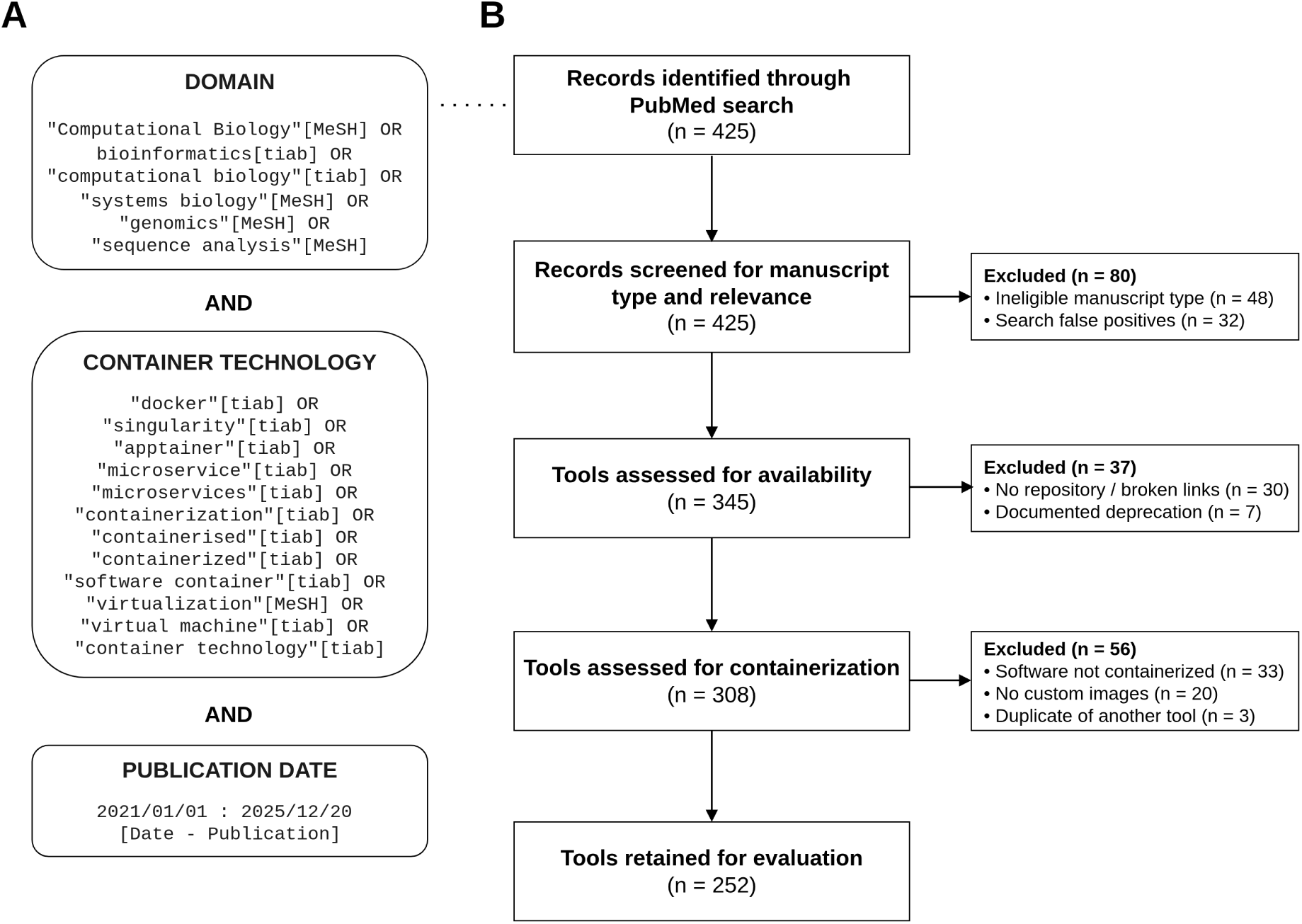
Tool review inclusion and exclusion results. PubMed query syntax (A) restricted search to containerized bioinformatics tools published between 2021-2025. The query returned 425 publications, of which 252 were included for final review after applying relevant filter criteria (B).

### Tool review and metadata

For each tool retained after the literature search and preliminary filtering, we compiled additional metadata relevant to containerization approaches and general software maintenance. At this stage, all characteristics were manually assessed based on the repository contents of the tool. Given that some tools included multiple component container images, we assessed all available container images to accommodate both tool-level and component-level analysis. We manually annotated the following metadata for all tools and components:

- **Image specification file path**: Relative path to specification file(s) to build the container image(s). Marked NA if none is present in the repository. The specification file is required to attempt to build the tool.
- **Built image repository and tag**: Repository and tag for any built image(s). Tags were annotated based on the most recently versioned tag; if none was available “:latest” was used. Marked NA if none was clearly specified in the paper or repository documentation. The built image allows us to confirm access and measure uncompressed image size after pulling locally.
- **License**: Software license specified for the given tool. Marked “unknown” if there was no license clearly provided in the tool repository, relevant documentation, or manuscript. Appropriate licensing is critical to distributing software, and this indicates how bioinformatics container developers are releasing tools and under what (if any) limitations.
- **Version control platform**: Version control platform for the tool (e.g., GitHub, GitLab, Bitbucket). The decision to leverage widely-adopted platforms can translate to more visibility and longevity of containerized tool source code.
- **Container engine**: Primary container engine used and/or documented. If multiple engines are mentioned they are combined as “;” delimited (e.g., Docker;Apptainer). Note that all references to Singularity are reconciled to the current name, Apptainer. The container engine gives context as to relative adoption of Docker and Apptainer in the community.
- **Orchestrator**: Container orchestration or execution platform described for the tool (e.g., Docker Compose, NextFlow). Note that for this category we consider workflow managers as container orchestrators. If no orchestration mechanism is clearly described, then the value is recorded as none. Tracking orchestration tooling being used will shed light on deployment needs and future solutions for making containerized bioinformatics tools more resilient.
- **Use case**: The primary use case for containerization for the given tool. Potential values are pipeline, web server/microservices, or standalone tool. Detailing how containers are used can help inspire adoption and inform future development of orchestration solutions.
- **Multi-arch**: Binary (TRUE or FALSE) as to whether there is evidence of developer documenting considerations and/or providing images compatible with multiple architectures (e.g., AMD64 and ARM64 builds published to DockerHub). Usage or mention of multi-arch techniques signals understanding of cross-platform compatibility challenges.
- **Multi-stage**: Binary (TRUE or FALSE) as to whether the container image uses multi-stage builds. If the tool includes multiple images, this is marked TRUE given at least one of the container image specification files uses a multi-stage build. Marked as unknown if no specification file is available. Multi-stage builds can cut down on image size and reduce the vulnerability surface. Evidence of adoption indicates community awareness and perception of benefits.
- **Parent image**: Tag for parent image(s) used in container image specification files. Marked as unknown if no specification file is available. This gives a picture of whether or not the community is building off of openly available, supported parent images. It also reveals practices for applying pinned tags for parent images or if developers tend to default to :latest tags.

In addition to the manually annotated metadata fields, we programmatically retrieved the full version control commit history for repositories, including date of first commit, date of last commit, and total number of commits. We used the GitHub API to infer the default branch and then cloned each repository to access the commit log for summarizing that branch. Version control data were retrieved in June 2026.

### Container image build and pull tests

One of the goals of our project was to assess whether each containerized tool could be rebuilt and/or pulled from its given repository. The scale of this review inspired us to conceive of an apparatus to automatically monitor and dispatch jobs to perform the image build and pull testing. We developed our platform, socr8s, as a containerized set of microservices that run via Kubernetes (K8S) orchestration. All socr8s components are specified as K8S resources, including a service to run a MongoDB container to retain logs, timing, and processing status as well as dispatcher services to monitor the job queues and launch build and pull jobs respectively. The stack also includes a ServiceAccount with RBAC details to allow the dispatchers to communicate directly with the K8S API. Under this configuration, when the build dispatcher sees a new tool marked as “queued” in MongoDB, it can launch the appropriate job pods to attempt the build based on the provided tool specifications in the database. The architecture is depicted in Supplementary Figure S1. Each build job is designed to run an image that includes Docker and a wrapper script to capture results (e.g., image size in bytes, failure or success, logs) and write output to the MongoDB database. The image pull dispatcher operates in the same fashion, but with a modified wrapper script to attempt a Docker pull command. The socr8s stack is available at https://github.com/vpnagraj/socr8s.

To conduct the build and pull tests for our study, we populated the socr8s infrastructure with the metadata curated during the initial literature review and screening. When uploading the metadata, we expanded tools with multiple component images to have one entry per component. The build and pull data was stored in separate MongoDB collections. All entries were initially marked with a “queued” status when uploaded. For the build procedure, we included a parameter for build context. In the initial upload, the context values were all set to the relative path for the image specification file. If the file was at the root of the repository, then the context would be recorded as “.”. No other instructions or steps for system-level customization were provided before the build attempt.

To run the socr8s components, we used a single-node Kubernetes cluster deployed with Minikube. The cluster was hosted on an Ubuntu 24.04 server and allocated 16 CPU Cores, 32GB RAM, and approximately 500GB of ephemeral storage. Pods were subject to constraints per resource specifications in the configuration YAML. The MongoDB was provided a minimum of 0.25 CPU and ∼0.25 GB RAM, with a ceiling of up to 1 CPU and ∼1 GB RAM available. The build and pull dispatcher pods were extremely lightweight, with a minimum 0.1 CPU and ∼0.125 GB RAM with up to 0.5 CPU and ∼0.25 GB RAM allowed. Build jobs were allocated a minimum 0.5 and up to 2 CPU, with at least ∼0.5GB and up to ∼4 GB RAM. Jobs initiated by the pull dispatcher were allocated exactly half the resources. The deployments were specified with a concurrency variable to constrain the number of jobs to no more than two build and pull jobs running simultaneously. Under this configuration, at any given time at most there were a total of four possible jobs, plus the MongoDB and the dispatcher services competing for resources on the cluster. There were no other pods running on the cluster during the build and pull tests. Build jobs were given a maximum of one hour to run before a timeout was recorded.

The socr8s apparatus was built to accommodate Docker images. For Apptainer tool testing, we chose to run *ad hoc* build scripts. We installed Apptainer version 1.5.0 and debootstrap 1.0 directly on the host machine and then scripted commands to retrieve the specification files, attempt a build, and record results to MongoDB.

The combination of manual review and ambiguity in build instructions (e.g., potential mismatches in build context expectations) motivated us to review and confirm any initial build failures detected. Expecting log output to be verbose, we looked for a way to leverage AI to assist in characterizing the failure state. Using Claude Opus 4.6, we built a skill to review container image build failures based on log output. The skill is available at https://github.com/vpnagraj/build-failure-review. For feasibility, we truncated log output to the last 1000 lines. We assessed the following failure categories: repository structure changed, context mismatch, dependency rot, vanished dependency, package conflict, parent image unavailable, resource exhaustion, network download failure, specification file parse error, and build script error. More extensive definitions of the categories used in the skill are available in Supplementary Table S1. In addition to the inferred failure modes, the build log review skill also suggests a level of difficulty (trivial, moderate, hard, or intractable) for resolving the issue. We assessed the precision of the build fix difficulty inferred by the AI via a *post hoc* experiment wherein we manually attempted a build directly on the host machine for all fixes deemed trivial.

Note that the build failure triage was the only circumstance in which AI was used for our study. We manually reviewed all failure categories ascertained by Claude and re-assigned failure mode in several cases. For any images where the failure audit indicated context mismatch, clear metadata entry inconsistencies (e.g., path to Dockerfile misspecified), or potential transient network drop out (e.g., Git clone of the tool repository in the builder pod failed unexpectedly), we removed the original failure entry from MongoDB and initiated a build retry with corrected information. We did not retry builds or attempt corrections for any other failures, and we restricted our failure audits to one retry.

Summary statistics are reported as median and interquartile range (IQR) for continuous variables. To analyze the build test results, we used odds ratio and 95% confidence interval as the measure of effect. All data retrieval, *ad hoc* MongoDB data updates (e.g., build retries), and analysis was conducted using R version 4.5.1 (R Core Team 2021).

## Results

### Literature search and tool screening

Figure 1 depicts the literature search approach, along with inclusion and exclusion results for the tools evaluated. The initial PubMed query returned n=425 articles, of which 252 were ultimately retained for subsequent tool evaluation. Of the initial search results, 48 articles were ineligible manuscript types (e.g., reviews, protocols, experimental studies that may have used containers but were not software papers). There were also 33 papers returned that described software that was not containerized. We excluded 3 papers that duplicated tools described in other search results. In addition, 32 results appeared to have been false positives that were not relevant to bioinformatics, often due to terminology overlap (e.g., singularity and Singularity). We found 37 tools for which there was no repository, broken links, or clearly documented deprecation notices. We also excluded 20 tools that used containers but did not appear to feature any custom images.

Table 1 summarizes baseline characteristics for the containerized tools. Among the n=252 software papers retained after initial screening, n=242 had links to source code in openly available version control repositories. About 93% of the repositories were available on GitHub, with the remainder either hosted on GitLab or Bitbucket. Historical development activity ranged from 1 to 5,064 commits, with a median of 131 (IQR: 37-349). About half had at least one commit within the two years leading up to our analysis.

**Table 1:** Summary of baseline characteristics of n=252 surveyed bioinformatics tools.

|  | n (%) |
| --- | --- |
| Version control platform |  |
| GitHub | 226 (89.7%) |
| GitLab | 10 (4.0%) |
| Bitbucket | 6 (2.4%) |
| None | 10 (4.0%) |
| Licensing |  |
| MIT | 86 (34.1%) |
| GPL | 71 (28.2%) |
| Other | 31 (12.3%) |
| Unknown | 64 (25.4%) |
| Use case |  |
| Standalone tool | 120 (47.6%) |
| Web server or microservices | 78 (31.0%) |
| Pipeline | 54 (21.4%) |
| Components |  |
| Single image | 209 (82.9%) |
| Multiple images | 43 (17.1%) |
| Container engine |  |
| Docker | 200 (79.4%) |
| Apptainer | 14 (5.6%) |
| Both | 38 (15.1%) |
| Orchestrator |  |
| None | 177 (70.2%) |
| Other | 16 (6.3%) |
| Docker Compose | 31 (12.3%) |
| Nextflow | 28 (11.1%) |
| Reviewable artifact |  |
| Both | 116 (46.0%) |
| Built image | 57 (22.6%) |
| Image specification | 79 (31.3%) |

When analyzing licenses applied, we found that approximately 1 in 4 tools did not specify a license. Among the licenses used, by far the most common were MIT (n=87) or some variant of GPL (n=71). Other types that appeared multiple times included versions of Creative Commons, BSD, Apache, Mozilla Public License. The remaining licenses were either custom or appeared in only one tool reviewed. The full distribution of licenses is available in Supplementary Table S2.

Roughly half of the tools (48%) were standalone software designed to execute a single bioinformatics task. The remainder were split between web server or microservices components (31%) and multi-step pipelines that use containers to encapsulate discrete tasks (21%).

For many tools we were able to review both a build specification and a pre-built image (46%), while the rest had only one or the other available. The large majority were distributed as a single container image (83%), with the remaining containing multiple component images, often aligning with the pipeline and microservices use cases noted above.

### Containerization practices

The distribution of container engines was highly imbalanced, with Docker predominating. Most tools that described using Apptainer did so alongside Docker by providing companion Dockerfiles, pre-built images in Docker container registries, or image conversion documentation. The majority of tools (70%) used no orchestrator at all, and among those that did, Docker Compose and Nextflow were the most commonly used mechanisms, which is again consistent with pipeline and microservice use cases.

We found that adoption of some containerization best practices was limited. Evidence of multi-architecture builds or documentation regarding platform compatibility was only present in about 7% of the tools with build instructions or architecture-specific tags in image registries. We found multi-stage build techniques used in only 10% of the tools with image specification files available. However, about 3 of every 4 tools used image tags that were pinned (i.e., not “:latest”), with many of the parents concentrating on a handful of types of images. About 39% of the tools were built on top of base R or Python images, with another 36% building on Debian or Ubuntu community images. Other less commonly used starting points included distributions of Fedora/CentOS and Alpine, as well as Node and OpenJDK images for Java tools. The full list is available in Supplementary Table S3.

### Image build evaluation

We found that the majority of tools with image specification files did not successfully build in our system. Even with retries to adjust parameters for context mismatch or transient network disconnections, only 26% were able to be built. For those that did, the built image size ranged between 0.01 and 27GB, with median 1.57GB (IQR: 0.72-2.93GB).

Table 2 summarizes the build success stratified by containerization practices. Of the practices we assessed, the strongest indicator that the image could build was recency of development activity. We found that tools with repositories that had at least one commit in the past two years were roughly twice as likely to build successfully (OR=2.26 (1.16-4.42)). The monthly version control activity for tools evaluated in the build test depicted in Figure 2 reinforces this result. For tools with non-ambiguous parent image tags, the OR point estimate leaned towards likelihood to fail, but the interval was wide and non-significant. When stratified by build success, multi-architecture considerations and multi-stage build practices had point estimates pointing in different directions. However, with only a handful of tools exhibiting either practice (n=14 multi-architecture; n=21 multi-stage), the confidence intervals for both measures were extremely wide and neither approached significance.

**Figure 2:**
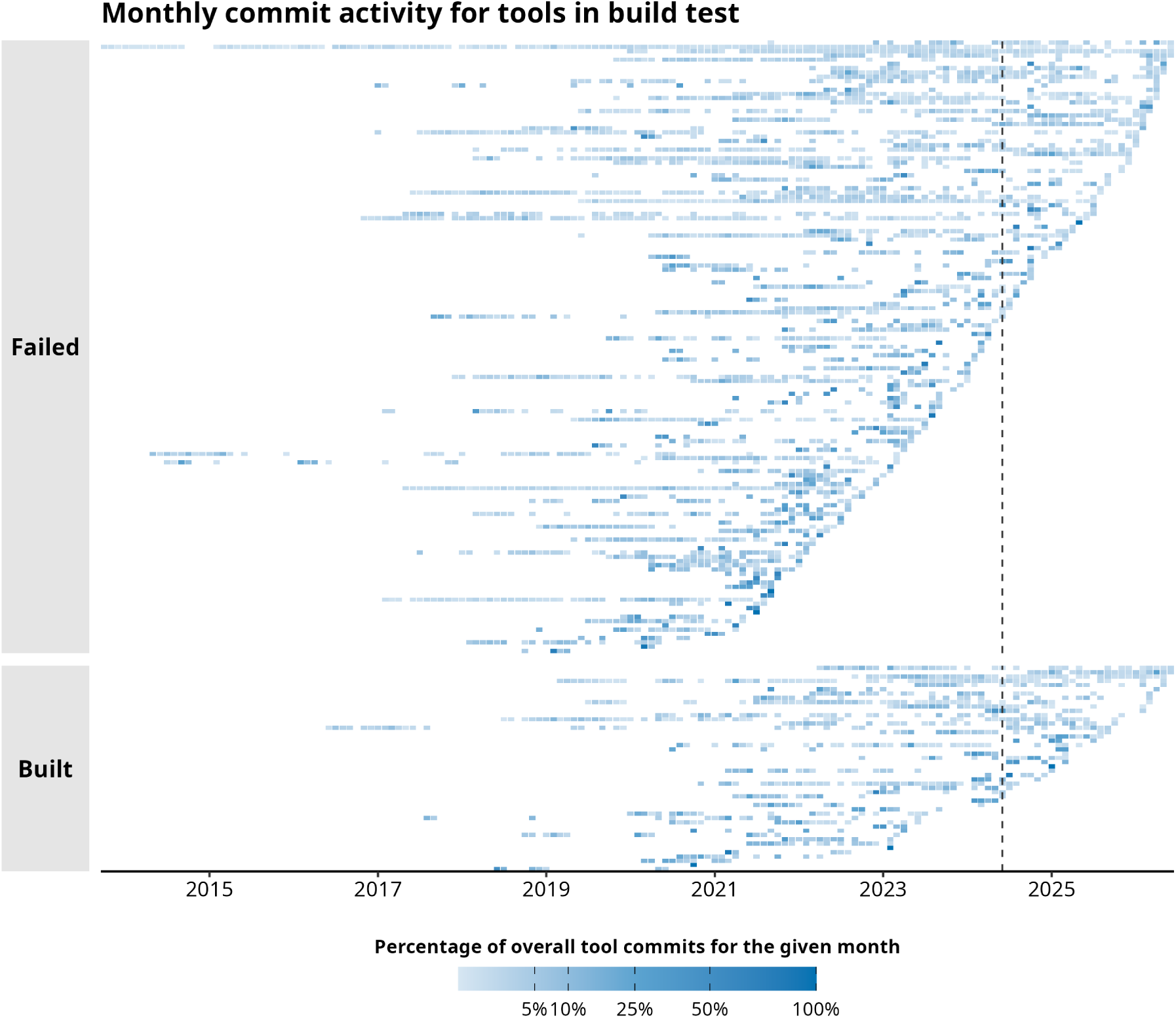
Commit activity for tools with image specification files. Each tool is represented as a row, with each tile representing a month of commit activity. Tile shading indicates percentage of overall commits for the tool that month. A darker shade indicates that the bulk of development occurred in the given month. Tools are stratified by build success. The vertical dashed line at 2024-06-01 marks the two years prior to the data retrieval for this analysis.

**Table 2:** Build success stratified by containerization practices for the n=195 tools with build specifications. For tools with multiple component images, build success status was only satistified if all component images could be built. The table includes an odds ratio of a successful build for the first level of each practice, relative to the reference level. Odds ratios that achieve statistical significance (p < 0.05) are marked with *.

|  | Build outcome |  |  |
| --- | --- | --- | --- |
|  | Successful (n = 50) | Failed (n = 145) | OR (95% CI) |
| Commit in last two years |  |  |  |
| Yes | 33 (66.0%) | 67 (46.2%) | 2.26 (1.16–4.42)* |
| No | 17 (34.0%) | 78 (53.8%) |  |
| Pinned parent image |  |  |  |
| Yes | 36 (72.0%) | 112 (77.2%) | 0.76 (0.37–1.57) |
| No | 14 (28.0%) | 33 (22.8%) |  |
| Multi-architecture build |  |  |  |
| Yes | 6 (12.0%) | 8 (5.5%) | 2.34 (0.77–7.10) |
| No | 44 (88.0%) | 137 (94.5%) |  |
| Multi-stage build |  |  |  |
| Yes | 4 (8.0%) | 17 (11.7%) | 0.65 (0.21–2.05) |
| No | 46 (92.0%) | 128 (88.3%) |  |

In addition to signals of containerization practices, we used logs to assess build failures. The AI-assisted log scan revealed that problems with dependencies or parent image availability accounted for about 30% of the build failures. The next most frequent class of failure was some form of error in the build script or image specification file syntax (27%), closely followed by an apparent change in repository structure (19%), which often manifested as files needed for COPY or ADD imperatives being unavailable. About 13% of the unsuccessful builds failed due to timeout or memory exhaustion. The remaining failures were due to lack of ability to authenticate to requested resources during build or network access.

Figure 3 presents failure categories for all n=145 tools that could not be built successfully. We stratified these counts by AI-determined difficulty for fixing the inferred failure cause. The vast majority (85%) of build failure fixes were deemed moderate or hard. Approximately 8% were characterized as having no feasible path to fixing the build failure. For example, some tools relied on parent images that were unavailable on open registries and lacked instructions to rebuild them in the given repository. On the other end of the spectrum, about 7% were considered trivial fixes. Some categories were notably divided between intractable and trivial fixes. For instance, the authentication required failures were generally split between inability to authenticate to assets (intractable) and terms of service consent required (trivial). We attempted our own *ad hoc* builds for the n=10 failures with fixes inferred to be trivial. We found that five could be fixed on a first attempt while the rest revealed secondary issues after resolving the initial error.

**Figure 3:**
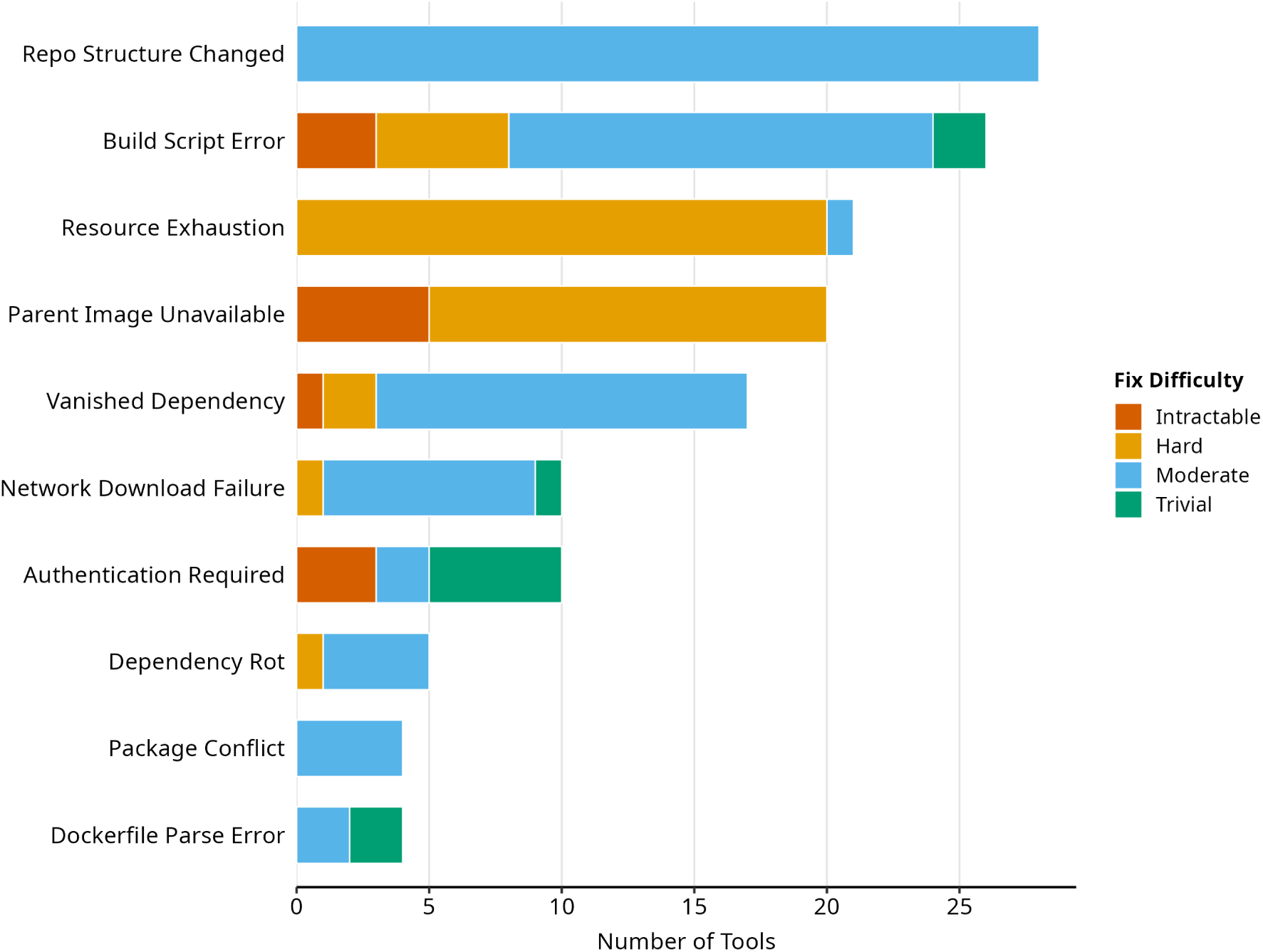
Count of build failures by inferred category and fix difficulty. Sum of bars equals n=145 tools that failed to build during testing.

### Image pull evaluation

Among the n=173 tools with images documented as being on a public container registry, nearly all could be pulled successfully. Only three images failed to pull, of which one had an OCI compliance issue on quay.io and the other two had outdated or broken registry links. As with the built images, pulled images tended to be large. Uncompressed size ranged between 0.01 and 88.75GB, with median 3.75 GB (IQR: 1.92-7.59GB). DockerHub was by the far the most common registry used, with n=8 tools hosting images on GitHub Container Registry (GHCR), n=4 on public GitLab registries, and n=3 on quay.io.

Note that after performing the image pull evaluation, we could assess overall ability to reconstitute or use the published containerized tool. Among the n=252 tools evaluated, n=61 (24%) failed to build and did not have pre-built images published in container registries.

## Discussion

### Findings

Software containerization has become a common and valuable open science practice in bioinformatics and many other disciplines. A container image specification gives developers and machines alike an explicit, executable account of a tool’s dependencies and expected runtime behavior. An image published to an open registry (e.g., DockerHub or GHCR) can often be pulled and run off-the-shelf for years after its release. But a container captures a tool at a single moment. As time moves on parent images may be deprecated, pinned dependencies might disappear, and host platforms and architectures will evolve. As our results show, the initial effort to containerize a bioinformatics tool is necessary but not sufficient to ensure that the tool remains accessible and functional over the long term. Containerization does not guarantee reproducibility. As an upfront practice, it must be paired with sustained maintenance to ensure that tools remain healthy and usable by others.

Our study highlights persistent gaps in the adoption of software development and sustainability practices among bioinformatics software developers. For example, approximately 25% of tools lacked a developer-defined license, creating unnecessary ambiguity around reuse, redistribution, and downstream integration. Although nearly all of the evaluated tools provided source code through public repositories, many exhibited declining contribution activity over time. Some degree of software deprecation is expected, particularly in rapidly evolving fields such as bioinformatics. However, responsible software stewardship requires clear documentation when projects are deprecated, archived, or no longer actively maintained. The combination of declining development activity and the limited number of tools explicitly designated as deprecated or orphaned suggests that many projects may be reaching end-of-life without clear communication of their maintenance status.

Researchers have previously documented a range of opportunities for improving bioinformatics development efforts (Russell et al. 2018), and our results situate those observations alongside contemporary containerization practice. On the whole, the tools we evaluated reflected an inclination towards published ideals for developing sustainable containerized software (Moreau and Wiebels 2026; Nüst et al. 2020). Where image specification files were available, the majority pinned parent tags rather than relying on moving targets upstream. More advanced techniques, however, were comparatively rare. Few tools took advantage of multi-stage builds, and both built and pulled images tended to be large (i.e., on the order of GB rather than MB). Large images carry practical consequences that users inherit in terms of transfer time, build cache, and storage overall. Orchestration beyond a single image was likewise rare. When orchestrators were used, they tended to be either domain-specific workflow managers (e.g., Nextflow) or lightweight tools (e.g., Docker Compose), while more contemporary architectures such as Kubernetes seldom appeared even in documentation. Although these patterns suggest a lag in the adoption of more advanced containerization technology, we read them less as a shortcoming than as a natural stage in the field’s maturation. Foundational practices have clearly taken hold, and there is a growing opportunity for community training and shared resources to help developers adopt the more advanced techniques that improve the efficiency, longevity, and portability of their tools.

One of the notable findings was that most tools with available specification files did not rebuild in our environment. We interpret this primarily as a consequence of software bitrot (i.e., drift towards breaking changes over time). Our build log review attributed about half of failures to broken dependency chains or shifts in repository structure that led to missing assets. After taking into account containerization practices, the strongest predictor of a successful build was recent development activity. Tools with a commit in the preceding two years were roughly twice as likely to build. Software that is not regularly maintained often lacks the tests, continuous integration, as well as other automated or manual touch points that would surface a broken build early. As failures accumulate over time, repair can become non-trivial. We found that the majority of failures required fixes of moderate or greater difficulty. Even seemingly simple fixes can belie greater challenges. For our *ad hoc* attempt to resolve failures with “trivial” fixes, only half resolved on the first attempt, while the remainder revealed secondary issues once the initial error was cleared. These findings reinforce how much ongoing attention a containerized tool requires to remain buildable.

The availability of pre-built images offsets some of these difficulties. Nearly all of the images located in public registries pulled successfully. In these cases, users have a reliable way to run a tool even when its specification file no longer builds. However, a registry copy is best positioned as a convenience for distributing tools, providing a safeguard rather than a guarantee of longevity. A pre-built image preserves a tool in the time and context it was built and pushed to the registry. To accommodate new features, security patches, or compatibility updates, the tool would need to be rebuilt. If the specification file or build instructions are no longer functional, there may be friction that delays or derails a release. It’s also worth emphasizing that in our analysis, 24% of tools were based on images that could neither be rebuilt nor pulled from a registry leaving no straightforward path to reconstituting the tool at all. For containerized software intended to remain useful over the long term, maintaining a build-able specification file is a worthwhile aim in its own right and should not be considered merely a secondary activity if built artifacts are published.

### Impact

Beyond providing specific findings, our study contributes a broad aperture and new methodology for studying containerized software sustainability. To our knowledge a focused survey of containerization practice at this scale (i.e., over 250 scientific computing tools systematically drawn from a single domain) has not previously been reported. Broader empirical work on Docker sustainability has already shown that containerization alone does not guarantee reproducibility in scientific computing (Malka, Zacchiroli, and Zimmermann 2026). Our study demonstrates that this pattern holds even within a single, tightly scoped domain. Large samples (e.g., hundreds of tools considered) make it possible to observe where real-world practice aligns with idealized recommendations, as well as where there is divergence. Our study does so at a scope that enables characterization of broad patterns, rather than individual anecdotes. The same data also speak to software stewardship considerations such as licensing and version control activity more generally. In this light, containerized tool health becomes recognizable as an important and understudied facet in how bioinformatics software is developed and sustained in general.

Methodologically, our design prioritized human expertise at multiple key points. We performed manual screening of search results, manual curation of tool metadata, and manual review of automated outputs, and did so at a level of granularity we believe is uncommon in surveys of this size. Where we did rely on AI, namely to triage build failure logs, we treated its classifications as a starting point and audited assignments to reclassify where warranted. Our experience with the failures flagged as “trivial” illustrates the merit of this approach. While we do not diminish the role that AI can play in making bioinformatics software more sustainable, we see our study as a practical example of how human judgment can and should be incorporated, particularly for activities that necessitate nuanced assessments of sustainability signals or software health.

Finally, our study yields artifacts that developers and users of containerized bioinformatics tools can immediately leverage. We have released the socr8s apparatus and the build failure review skill so that the buildability of a containerized tool can be more reliably assessed rather than discovered *ad hoc*. For maintainers, such an assessment can flag latent fragility early and point toward concrete steps to make a tool more robust and dependable. For researchers who depend on a tool downstream, a triaged build failure can offer a structured account of what went wrong, which can shorten the path to restoring a tool to a functional state or adapting it to support new features. Our expectation is that lowering the effort required to evaluate and repair containerized software makes ongoing maintenance a more approachable undertaking rather than an open-ended burden.

### Limitations

We acknowledge that our study has several limitations. While we see human judgement as a key strength of our study design, manual assessment inevitably introduces a degree of subjectivity. It is possible that another team applying the same criteria could arrive at slightly different annotations. We mitigated this by defining explicit criteria for each metadata field in advance, and, where possible, pairing human review with automation and AI-generated classifications. Our guidance in key steps of building the AI review skill (e.g., refining failure categories) was subjective. However, delivering the skill as a versioned artifact makes these decisions auditable and open to revision, allowing consensus to build over time with future contributions. Ultimately, we see the manual component of the study not as a source of unchecked human bias but as expert judgment exercised within a structured framework.

Our measurement of development activity contributes important context for our analysis. However, using commit history for the entire repository rather than for the container specification file alone is an imperfect proxy for attention paid specifically to containerization. Some commits we counted will have been unrelated to the image. It nonetheless serves as a reasonable signal of whether a tool is being actively maintained, particularly because our sample was restricted to tools published in dedicated software papers and many container images were delivered as standalone tools.

The build apparatus was deliberately automated and standardized, and it was not designed to accommodate bespoke configurations, required build steps documented only in prose, or non-standard wrappers (e.g., invoking docker build through a Makefile). Some tools that failed in our system might build successfully under hands-on, tool-specific intervention. This constraint is intentional and informative insofar as it approximates the experience of a non-expert user who discovers a tool and attempts a build. In this scenario, a user could conceivably navigate a bespoke build process, but this is unlikely and misses some of the core advantages of standardization that containerization offers. Also related to the build system, our choice to fix resource constraints ensured comparability across build tests but results could differ under other resource allocations or network configurations. We concede that a small number of timeout or resource-exhaustion failures might resolve given more generous limits.

We restricted our study to tools that were published in peer-reviewed software papers, and all initial literature was drawn from a single bibliographic source. While PubMed is the natural repository to look for bioinformatics literature, our methods may have missed outputs that were indexed elsewhere or not peer reviewed. Our findings should accordingly be read as describing tools accompanied by a software publication rather than a full accounting of all containerized bioinformatics software.

Lastly, our study at best captures a single moment in time. The specific tools we evaluated will continue to evolve, with build failures being repaired, images updated, and repositories archived after our analysis window. We cannot claim that these results predict future practice or developer attitudes. New technologies, including generative AI, may become embedded in the workflows and environments bioinformatics researchers use, potentially in ways that strengthen containerization and software development practices over time. However, capturing a baseline is required for measuring whether and how such practices change and we intend our findings to serve as a reference point for that prospective view.

### Future directions

Containerization is a powerful reproducible scientific computing technique. This is especially true for domains like bioinformatics, where software often depends on complex and evolving stacks of system libraries, languages, and community maintained packages. Our study provides a baseline view of current trends in real-world bioinformatics containerization techniques. Future work to track the evolution of these practices as they evolve will be critical, especially as the container technologies advance and adoption of generative AI pushes software development in new directions. We anticipate that containerized tools will continue to proliferate, and applying sound practices upfront may be more achievable with AI uplift. While the initial attention to practices like avoiding image bloat and appropriately pinning dependencies is critical, without ongoing care even well-built images can drift towards the bitrot we observed. Longitudinal study of how containerization practice and tool health change over time would help distinguish durable improvement from fleeting polish.

Concerted efforts could move the field toward wider and more effective usage of containerized tools. Emphasis on training and upskilling can broaden familiarity with both advanced containerization techniques and general maintenance pitfalls. Developer-facing solutions similar to those we released for this study could make it more feasible to audit tool quality and ideally steer maintainers toward sustainable practices. As tools continue to be published, journal editors and reviewers could play a vital role in identifying sustainability signals in software papers. There is a growing opportunity for community guidelines that more concretely connect to practical behaviors and help users, reviewers, and the maintainers move beyond confirming that a container image is simply described toward looking for the markers of health and sustainability.

While our focus has been on bioinformatics, the phenomena we describe are not unique to it. Containerization increasingly underpins reproducibility efforts in genomics-adjacent biomedical fields, the physical sciences, and computational research broadly. The tension between upfront concessions made to expedite initial build and the sustained effort required to keep software alive is common in other domains. We expect that evaluation in more diverse disciplines could help inspire broader and more confident usage of container technology for scientific computing applications at large.

