## Supplementary Material for "Survey and Evaluation of Applied Containerization Practices in Bioinformatics"

VP Nagraj Stephen D Turner Neal Magee

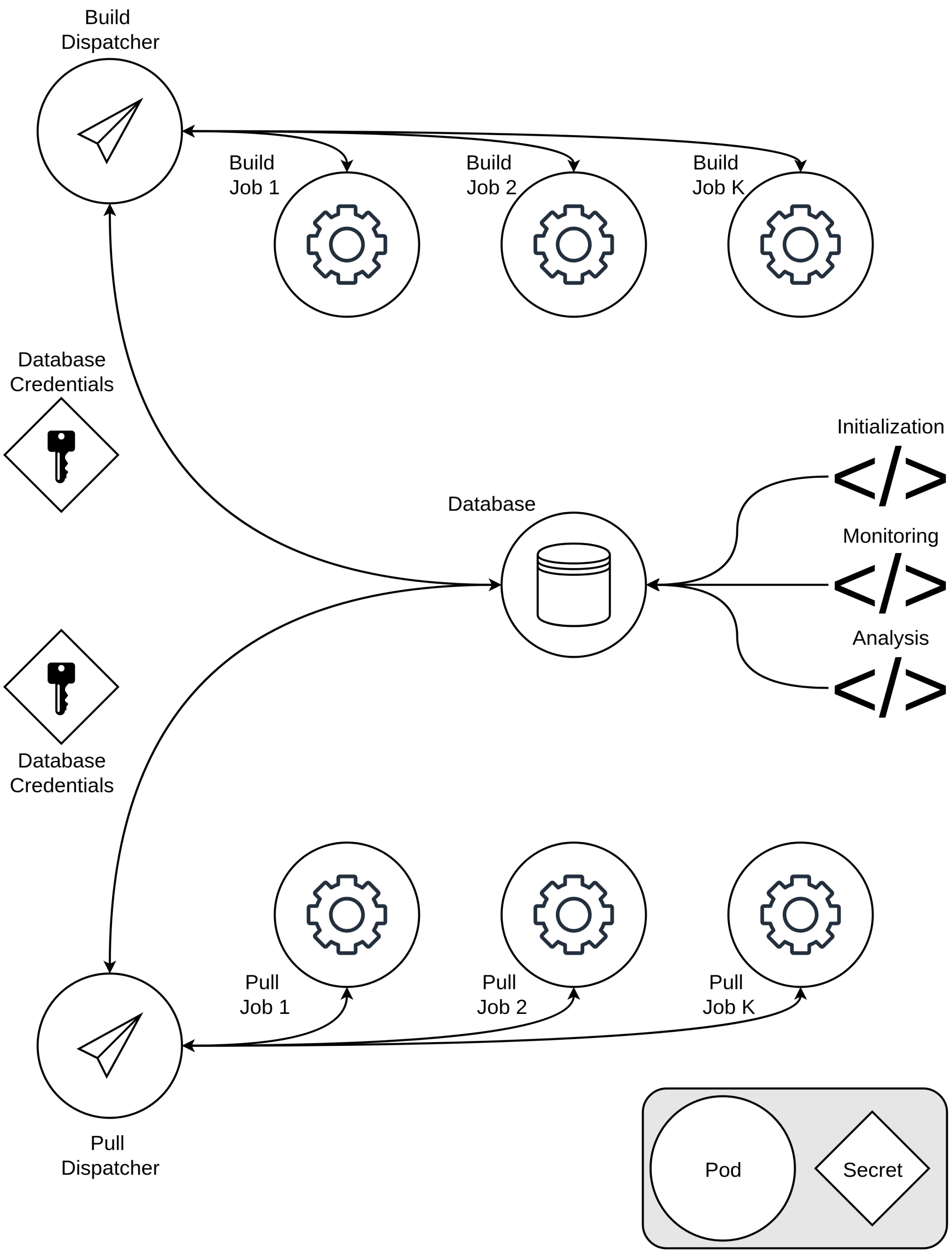


Figure S1: Illustration of the socr8s build and pull apparatus. The system uses a collection of Kubernetes resources to manage a queue for jobs to be dispatched automatically. The build and pull dispatchers communicate directly with the MongoDB database to look for queued entries. The dispatcher pods have access to the Kubernetes API and launch jobs with specifications based on the given database entry. Each job is passed secrets to authenticate to the database and submit logs. The database service can be exposed to enable programmatic access from outside of Kubernetes for queueing, monitoring, and analyzing job output.

Table S1: Container image build failure taxonomy used by the build log review AI skill. The taxonomy was generated by Claude and then iteratively refined with human review.

| Category | Description |
|---|---|
| **Dependency Rot** | Parent image's package manager or tooling is too old to talk to current upstream repos. The tools exist but can't resolve/download. |
| **Vanished Dependency** | A specific pinned package version, download URL, or release artifact no longer exists upstream. |
| **Parent Image Unavailable** | The FROM image exists but its underlying OS repos have been decommissioned (e.g. Debian stretch, Ubuntu old releases). This is "vanished" at the distro level. Keep it separate from Vanished Dependency because the fix is a full rebase, not a pin change. |
| **Network Download Failure** | A download hangs, times out, or is rate-limited. The resource may or may not still exist. Includes FTP timeouts and transient HTTP errors. |
| **Context Mismatch** | The Dockerfile COPYs or ADDs files that exist in the repo but are outside the Docker build context directory. |
| **Repo Structure Changed** | The Dockerfile references files, paths, or scripts that no longer exist in the current state of the repo. |
| **Auth Required** | The build tries to fetch from, or operate against, a gated source. Covers three sub-cases: (a) credential/login walls, (b) license- or registration-gated downloads, and (c) terms-of-service assent that cannot be given in a non-interactive build. Conda's TOS gate is the canonical case. |
| **Resource Exhaustion** | The build ran out of memory, disk, or exceeded a time limit during compilation or package installation. |
| **Platform Incompatibility** | The Dockerfile assumes a specific architecture, GPU, or OS feature not available in the build environment. |
| **Dockerfile Parse Error** | Docker cannot parse the Dockerfile at all. The build fails before any layer runs. Strictly pre-execution: bad instruction syntax, malformed heredoc, unknown directive, invalid FROM. If a RUN command actually executed and then failed, it is NOT this category. |
| **Package Conflict** | Packages from different eras or sources conflict with each other during installation. The conflicting packages all exist and download fine, they just can't coexist. dpkg/rpm file overwrites, mutually incompatible version constraints. |
| **Build Script Error** | A command or script executed during the build fails for reasons intrinsic to the code or build logic (e.g., compilation errors, missing headers, bad flags, a stricter compiler, or a logic error in a RUN line). Classify here only after ruling out the dependency/network/resource/platform categories above, since those also surface at execution time. |

Table S2: Full distribution of licenses observed in evaluated tools. Note that if a tool indicated multiple licenses in a repository or documentation, all were retained and separated by ';'. Tools without licensing clearly specified were characterized as having 'UNKNOWN' licenses.

| License | N |
|---|---|
| MIT | 86 |
| UNKNOWN | 64 |
| GPL-3 | 55 |
| Apache | 7 |
| BSD-3 | 7 |
| AGPL | 6 |
| GPL-2 | 6 |
| Custom | 5 |
| Custom non-commercial | 2 |
| Mozilla Public License | 2 |
| BSD-2 | 1 |
| BSD-3; CC0 | 1 |
| CC-BY-SA-NC | 1 |
| CC-NC | 1 |
| CC-SA | 1 |
| CC0 | 1 |
| CeCILL | 1 |
| EUPL | 1 |
| GPL | 1 |
| GPL-3; MIT | 1 |
| LGPL-2 | 1 |
| LGPL-3 | 1 |

Table S3: Unique parent images across all n=222 components of n=195 unique tools with image specification files available. Results are binned by the most widely used types of parent images, with all others aggregated to 'Other'.

| Parent flavor | Unique parent images (count) |
|---|---|
| R | bioconductor/bioconductor__docker (1); bioconductor/bioconductor__docker:3__21 (1); bioconductor/bioconductor__docker:RELEASE__3__13 (1); bioconductor/bioconductor__docker:RELEASE__3__17 (1); bioconductor/bioconductor__docker:RELEASE__3__18 (1); bioconductor/bioconductor__full:RELEASE__3__10 (1); r-base (1); r-base:3.4.3 (1); r-base:4.0.2 (1); r-base:4.0.3 (1); r-base:4.0.4 (1); r-base:4.3.2 (1); rocker/r-base (1); rocker/r-base:4.1.0 (1); rocker/r-ubuntu:20.04 (1); rocker/r-ver:3.5.0 (1); rocker/r-ver:3.6.3 (1); rocker/r-ver:4 (1); rocker/r-ver:4.0.0 (1); rocker/rstudio:3.6.3 (1); rocker/rstudio:4.5.0 (1); rocker/rstudio:latest (1); rocker/shiny (1); rocker/shiny-verse:3.6.1 (1); rocker/shiny-verse:4.2.3 (1); rocker/shiny-verse:latest (1); rocker/shiny:3.6.2 (1); rocker/shiny:4.2.1 (1); rocker/shiny:4.3.3 (1); rocker/tidyverse:4.4.0 (1) |
| Python | continuumio/miniconda3 (9); python:3.10 (3); python:3.10-bullseye (3); conda/miniconda3 (2); python:3.10-slim (2); python:3.12-slim (2); python:3.12-slim-bookworm (2); python:3.8 (2); condaforge/mambaforge (1); condaforge/mambaforge:latest (1); condaforge/miniforge3:24.9.2-0 (1); continuumio/miniconda3:23.10.0-1 (1); continuumio/miniconda3:25.3.1-1 (1); continuumio/miniconda3:4.10.3 (1); continuumio/miniconda3:4.7.12 (1); mambaorg/micromamba (1); mambaorg/micromamba:0.25.0 (1); mambaorg/micromamba:0.25.1 (1); mambaorg/micromamba:1.3.1 (1); mambaorg/micromamba:latest (1); python:2.7.10 (1); python:2.7.14 (1); python:3-slim (1); python:3.11-bullseye (1); python:3.11-buster (1); python:3.11.13-slim-bullseye (1); python:3.12 (1); python:3.12-slim-bullseye (1); python:3.12.7 (1); python:3.13-slim-bullseye (1); python:3.7-slim-bookworm (1); python:3.7.9 (1); python:3.7.9-slim-buster (1); python:3.8-bullseye (1); python:3.8-slim-buster (1); python:3.9 (1) |
| Ubuntu/Debian | ubuntu:18.04 (18); ubuntu:20.04 (17); ubuntu:22.04 (13); ubuntu:16.04 (7); ubuntu:latest (6); ubuntu:focal (2); debian:11 (1); debian:9-slim (1); debian:bookworm (1); debian:bookworm-slim (1); debian:bullseye-slim (1); debian:jessie (1); debian:stable-slim (1); ubuntu:14.04 (1); ubuntu:19.10 (1); ubuntu:21.10 (1); ubuntu:sha256.80c52afadf3e7c3f9573de4fe79b7dca57fb3290df6c8dc46a75b02768a81146 (1) |
| Other | amancevice/pandas:1.4.0-slim (2); debootstrap (2); docker.io/maven:3-eclipse-temurin-21 (2); libddocker/ubuntu16.04__base:latest (2); node:16-alpine (2); quay.io/bgruening/galaxy:20.09 (2); Clomet__v0/Dockerfile (1); adoptopenjdk/openjdk11:jdk-11.0.6__10-slim (1); adoptopenjdk/openjdk11:jre-11.0.8__10-alpine (1); adoptopenjdk/openjdk13 (1); aklimate-tmp (1); alpine:3.12.0 (1); alpine:3.13 (1); alpine:3.7 (1); amancevice/pandas (1); amancevice/pandas:1.0.1 (1); amazoncorretto:21 (1); amd64/ubuntu:24.04 (1); biobakery/mmuphin (1); biocorecrg/debian-perlbrew:buster (1); brcloudproject/ubuntu18.04-mpich (1); broadinstitute/gatk:4.1.4.0 (1); broadinstitute/gatk:4.5.0.0 (1); cchon/scafe:v0.9.0-beta (1); centos:7 (1); centos:7.6.1810 (1); ddecap/halvade-base:latest (1); docker.artifactory.pnnl.gov/pspecter/pspecterbase:2.0.0 (1); docker.io/alpine:3.21.3 (1); docker.io/ensemblorg/ensembl-vep:release__111.0 (1); docker.io/maven:3-openjdk-11 (1); docker.io/openresty/openresty:1.21.4.1-7-alpine-fat (1); eclipse-temurin:11-jdk-jammy (1); eclipse-temurin:11-jre-jammy (1); etal/cnvkit:0.9.7 (1); fedora:33 (1); fedora:35 (1); gcr.io/distroless/java11-debian11 (1); gcr.io/distroless/java21-debian12 (1); ghcr.io/hugheylab/stocker:latest (1); ghcr.io/marcelauliano/mitohifi-base:b3.2.3 (1); ghcr.io/pm4onco/cbioportal:latest (1); ghcr.io/rothamsted/knetminer-bare:latest (1); ghcr.io/stanstrup/qc4metabolomics/qc__base:latest (1); giovtorres/docker-centos7-slurm:17.02.9 (1); golang:1.13.7-buster (1); golang:1.17.2-alpine (1); http://mirror.centos.org/centos-8/8/BaseOS/x86__64/os/ (1); jboss/wildfly (1); jupyter/scipy-notebook:abdb27a6dfbb (1); koki/tensorlycv__component:latest (1); libd/r3.4.3__base:latest (1); libddocker/r__3.6.1__bioc (1); libddocker/samtools-1.3.1:latest (1); libddocker/samtools:1.10 (1); libddocker/samtools:1.9 (1); lifebitai/shapeit4 (1); mariadb (1); mariadb:10.8.2 (1); mariadb:11.3.2 (1); maven:3.6.3-openjdk-15 (1); mcr.microsoft.com/dotnet/runtime:8.0-alpin (1); mysql:debian (1); ncbi/blast:2.14.0 (1); nfcore/base:1.13.3 (1); nfcore/base:1.9 (1); nfcore/base:latest (1); nginx (1); nginx:1.17-alpine (1); nginx:alpine (1); nginx:stable-alpine (1); node:13.12.0-alpine (1); node:14 (1); node:18-alpine (1); node:18.16.1-alpine (1); node:20-buster (1); node:20.12.2-alpine (1); node:24-alpine (1); node:lts-alpine (1); nvidia/cuda:10.1-devel-ubuntu18.04 (1); nvidia/cuda:11.0-cudnn8-runtime-ubuntu18.04 (1); nvidia/cuda:11.8.0-base-ubuntu20.04 (1); omatt/default/mpi:3.1.1 (1); omicsnotebook-base (1); openjdk:11-jdk (1); openjdk:15-alpine (1); openjdk:15-jdk-alpine (1); openjdk:8u212-jdk (1); parkinsonlab/architect:latest (1); pegi3s/cutadapt:latest (1); pegi3s/docker:29.0.1 (1); pkuksa/bts-base:latest (1); postgrest/postgrest:v9.0.1 (1); quay.io/bgruening/galaxy:20.05 (1); quay.io/biocontainers/samtools:1.15–h1170115__1 (1); quay.io/broadinstitute/viral-baseimage:0.1.20 (1); quay.io/cellgeni/jupyter:r-base-20210222 (1); quay.io/qiime2/amplicon:2024.10 (1); redis:latest (1); ribocleaner-core:latest (1); rnakato/custardpy__r:3.4.1 (1); rnakato/mapping:2026.03 (1); rnakato/mapping:2026.04 (1); rnakato/r__python__gpu:2026.03.2 (1); rnakato/shortcake__seurat:3.5.0 (1); ruby:3.0.0-slim (1); rust:1.69-bookworm (1); rust:1.72-bullseye (1); sherlockdatalake/base:8u212 (1); tensorflow/tensorflow:2.8.2-gpu (1); tomcat:8.5.41-jre8-slim (1); tomcat:9.0.58-jdk17-openjdk-slim (1); tomcat:9.0.65-jre8 (1); trinityctat/starfusion:1.9.0 (1); vanallenlab/miniconda:3.12 (1); vibpsb/i-adhore:3.1 (1); xiaoxiutan/pgnneo:v1 (1); yum (1); zhai2018/galaxy:19.05 (1) |
